# A Detailed Map of Coupled Circadian Clock and Cell Cycle with Qualitative Dynamics Validation

**DOI:** 10.1101/2020.11.17.386110

**Authors:** Adrien Rougny, Loïc Paulevé, Michèle Teboul, Franck Delaunay

## Abstract

The temporal coordination of biological processes by the circadian clock is an important mechanism, and its disruption has negative health outcomes, including cancer. Experimental and theoretical evidence suggests that the oscillators driving the circadian clock and the cell cycle are coupled through phase locking. We present a detailed and documented SBGN PD map of known mechanisms related to the regulation of the circadian clock, and its coupling with an existing cell cycle SBGN PD map which includes main interactions of the mammalian cell cycle. The coherence of the merged map has been validated with a qualitative dynamics analysis. We verified that the coupled circadian clock and cell cycle maps allow reproducing the observed sequence of phase markers. Moreover, we predicted mutations that contribute to regulating checkpoints of the two oscillators. Strikingly, our approach underlines the potential key role of the core clock protein NR1D1 in regulating cell cycle progression. We predicted that its activity influences negatively the progression of the cell cycle from phase G2 to M. It is consistent with the earlier experimental finding that pharmacological activation of NR1D1 inhibits tumour cell proliferation and shows that our approach can identify biologically relevant species in the context of large and complex networks.

## 1 Introduction

The circadian clock is an adaptive mechanism that allows most organisms to anticipate the daily variations in their environment resulting from the rotation of the earth. In mammals, this system is present in virtually all cells and consists in a complex genetic network including interconnected transcriptional/translational feedback loops and that oscillates with a period close to 24h. This mechanism is interacting with many other pathways to temporally coordinate basic cellular functions such as signalling, metabolism, transport, autophagy and the cell cycle. Although circadian rhythms of cell division have been observed for nearly a century, it is only recently that molecular mechanisms connecting the circadian clock and the cell cycle mechanisms have been uncovered [29, 19, 21]. Importantly, the dynamics of the two systems were shown to be robustly coupled in dividing mammalian cells [14, 6]. This coupling suggests bidirectional interactions, however, while several mechanisms explaining the action of the circadian clock on the cell cycle have been proposed, how the cell cycle may influence the clock dynamics remains unclear. This issue is of primary conceptual and biomedical significance because how cell integrate multiple oscillators remains to be understood while accumulating evidence supports the notion that circadian disruption is a hallmark of cancer cells in addition to deregulated cell cycle, apoptosis and DNA repair pathways, avoiding immune destruction, metabolic reprogramming, etc. [12]. A better understanding of these links may also pave the way to the development of timed chemotherapy [3].

Given the complexity of the cell cycle machinery, addressing experimentally this issue is particularly challenging, this being mainly due to the fact that perturbation experiments are not specific and often difficult to interpret as a result of numerous, complex, or even artifactious interactions. Mathematical modelling has become instrumental as a complementary approach to the classical genetics and biochemical analysis of complex dynamical biological systems such as the circadian clock and the cell cycle [11]. In contrast to classical continuous or stochastic modelling frameworks which remain limited to a relatively low number of variables, qualitative and logical models are able to accommodate large networks [35, 38]. The availability of a detailed Systems Biology Graphical Notation Process Description (SBGN PD) map of the cell cycle including all key molecular mechanisms underlying its regulation prompted us to build a circadian clock counterpart.

The SBGN PD map we introduce provides a static representation of known molecular mechanisms related to the regulation of the circadian clock. It is designed as to also include a number of species making the interface between both cycles and playing a key role in their coupling. In order to check the validity of both maps with respect to the coupling of the two cycles, we employ computational methods to merge them into a larger map and build a dynamical model from it. We subsequently verify that our model can reproduce essential known behaviors of each cycle or of their coupling. Aiming at introducing as few *a priori* as possible, notably related to kinetic parameters and molecular concentrations, we rely on a qualitative modelling approach. In this approach, we consider that a molecule or complex is either absent or present, and compute their possible state changes according to the reactions specified in the map. Being coarse grain with very few parameters, qualitative analysis allows reasoning on large networks and providing robust predictions [32].

We show that our qualitative dynamical model can reproduce the observed sequences of activation of markers during the circadian clock and cell cycle progression. Our qualitative analysis also enables verifying and predicting checkpoints, i.e., steps or molecular activities that are necessary for the progression of the cycles. Those predictions can uncover key interactions between the coupled cycles, by showing how one cycle can temper with the progression of the other cycle by regulating part of its checkpoints.

## 2 Results

### 2.1 The circadian clock map

Several maps of the circadian clock mechanism have been generated and made publicly available through the PANTHER (http://www.pantherdb.org/), Reactome (https://reactome.org/) or KEGG (https://www.genome.jp/kegg/) databases. Although these interactive maps are useful tools for data exploration and pathway analysis, they all suffer from specific limitations ranging from absence or over abstraction of key regulatory modules to unbalanced complexity, to format conversion and layout rendering issues. This makes their direct use as sub-networks of larger maps tedious, time consuming if not virtually impossible. We therefore undertook the construction a novel map of the mammalian core circadian clock devoid of such limits with two main objectives in mind: 1) the map should be able to capture the known dynamics of the circadian clock when analyzed using the qualitative modelling approach used in this paper and 2) allow the merging with an existing SBGN PD map recapitulating the cell cycle network [10]. The SBGN PD circadian clock map reported here displays key experimentally validated nodes and their interactions known to play a significant role in the dynamics of the core mammalian circadian clock (Fig. 1). It contains 68 species including 14 genes, 14 mRNA and 47 proteins (represented by 154 different chemical species), 4 simple molecules and heat. One hundred and sixteen reactions include 21 associations, 7 dissociations, 28 transcriptional and translational regulations, 40 post-translational modifications and 20 transports occurring in either the cytoplasm, nucleus or extracellular compartments. Heat and glucocorticoids (GC) displayed in the extracellular space represent important inputs contributing to the synchronization of peripheral clocks [7, 4]. The circuitry shown in this map was assembled from several biologically meaningful modules. For instance, the CLOCK:BMAL1 heterodimer and its associated posttranslational regulation by PARP, CK2*α*, GSK3*β*, and SIRT1 forms the core of the nuclear module controlling the transactivation of 8 different targets (*Nr1d1*, *RORc*, *Cry1*, *Cry2*, *Per1*, *Per2*, *Wee1*). As NR1D1 undergoes extensive posttranslational regulation by GSK3b, HUWE1, MYCBP2, and FBXW7, which is potentially important in the context of the coupling with the cell cycle, this was detailed in another module. The PER et CRY proteins form the core of both a cytoplasmic module that describes their association and posttranslational regulation by kinases (CK1d, CK1e), ubiquitin ligases (*β*TRCP, FBXL3) deubiquitinases (USP2, HAUSP) and a nuclear module recapitulating the mechanism through which they inhibit the transcriptional activity of the CLOCK:BMAL1 heterodimer. This module is connected to the p53 nuclear module through PER2. Finally, a module recapitulating the action of HSF1 and glucocorticoid signaling on *Per2* gene expression was included as synchronizing inputs. Six proteins present within the cell cycle map and connected to the clock network through physical interaction or gene regulation have been included in the circadian clock map. These proteins act at various phases of the cell cycle and the directionality of the influence is primarily from the clock to the cell cycle with only 2 interactions out of 8 being in the opposite direction (Table 1). Although not present in the cell cycle map the MYC protein and its inhibitor MIZ1 have been included as this link may be relevant to the cancer cell context [41]. Upon visualization in CellDesigner, non experts can easily navigate this map which is further augmented by nearly 300 links to external databases in the present version.

**Figure 1:**
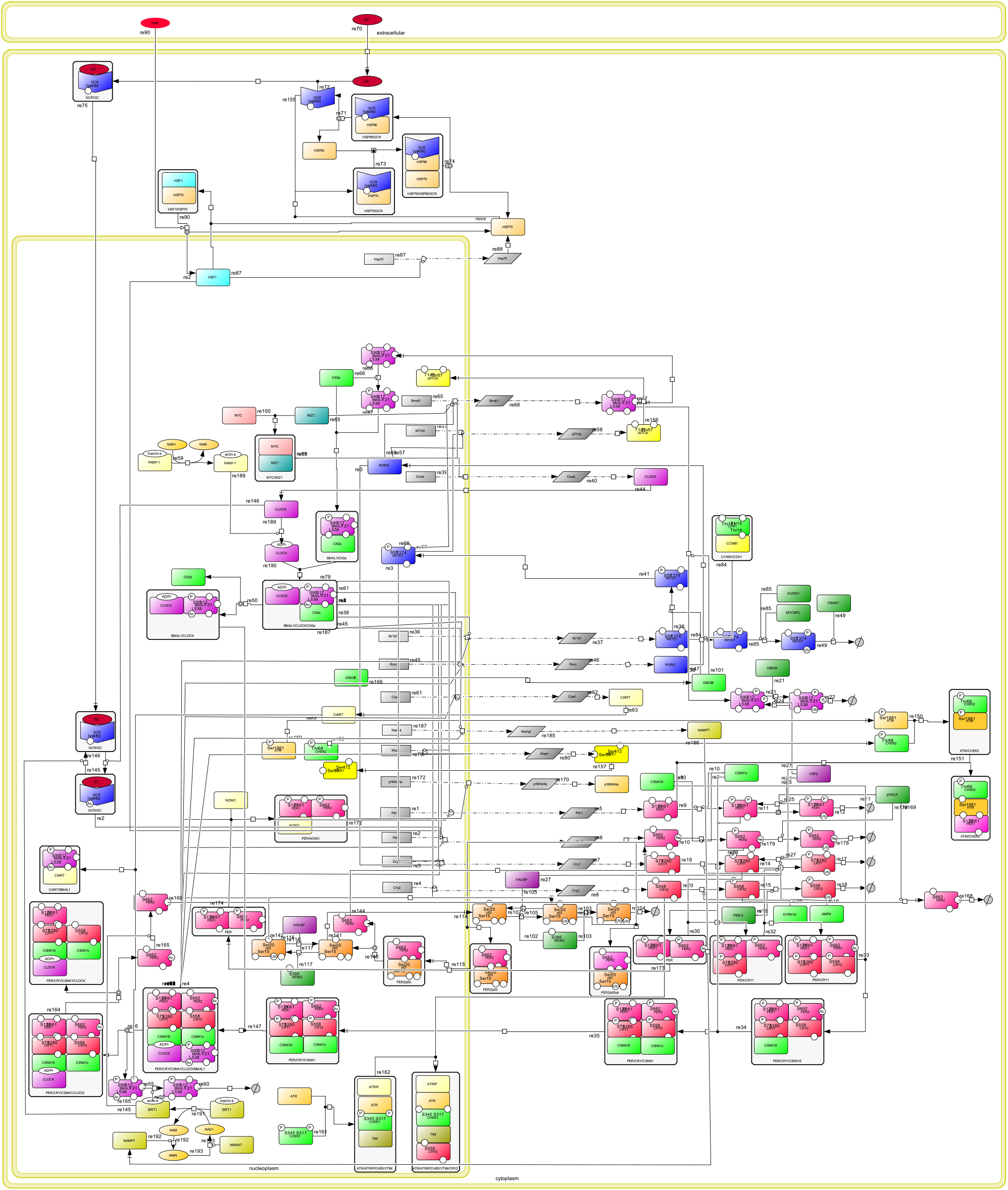
Overview of the circadian clock map, visualized in CellDesigner. This map shows the main biomolecular processes underlying the circadian clock. It is represented using the CellDesigner format [15], which is compatible with the SBGN Process Description language [40].

**Table 1:** Summary of experimentally validated molecular interactions between the circadian clock and the mammalian cell cycle molecular networks. Arrows represent the direction of the influences (→: from the circadian clock to cell cycle; ←: from the cell cycle to the circadian clock). All reported interactions have evidence in the literature.

| Clock |  | Cell Cycle | Phase | Mechanism | Output | Ref. |
| --- | --- | --- | --- | --- | --- | --- |
| BMAL1:CLOCK | → | *Wee1* | G2/M | transcriptional | activation | [29] |
| NR1D1 | → | *p21* | G1 | transcriptional | repression | [21] |
| RORc | → | *p21* | G1 | transcriptional | activation | [21] |
| NONO | → | *p16/Ink4a* | G1 | transcriptional | activation | [25] |
| NR1D1 | ← | CDK1 | G2/M | phosphorylation | degradation | [47] |
| PER1 | → | CHEK2/ATM | S | complex formation | DNA damage induced apoptosis | [19] |
| CRY2/TIM | → | CHEK2/ATM | S | complex formation | defective intra-S checkpoint | [45] |
| PER2 | ← | p53 | S | nuclear translocation | stabilisation | [20] |

### 2.2 Merging of the circadian clock and cell cycle maps

We built a map of the coupling of the circadian clock and the cell cycle by merging our circadian clock map and the cell cycle map into a larger map. The merging first required adjusting the specification of all entities shared by both maps for unifying their names and their list of modified residues. Based on our knowledge of the two maps, we therefore manually listed these entities, and the required modifications. Rather than proceeding to these modifications manually, we then implemented them in a program using the *csbgnpy* Python library, which allows manipulating the conceptual models underlying SBGN PD maps (see Materials and Methods for more details). Such a programmatic approach has several advantages: it allows using the original maps as inputs, it makes the list of modifications explicit, and the merging can be repeated on updated versions of the input maps. All changes made to both maps are described in details in Supplementary Information section 1.

### 2.3 Validation using qualitative dynamical analysis

In order to validate the content of both maps with respect to the coupling, we built a qualitative dynamical model from the merged map, and checked whether our model could reproduce known behaviors of both cycles and their coupling. Specifically, we checked whether our model could reproduce a correct progression of both cycles, and predict correct influences of entities of one cycle on the progression of the other.

The progression of each cycle is characterized by a (repeated) succession of phases. Hence, for each cycle, we checked whether our model could reproduce a correct succession of these phases. To achieve that, we considered a number of markers for each phase of both cycles, that are entities specifically present in that phase, and checked whether they could be reached from a specific initial state under different conditions (see Fig. 2). The cell cycles exhibits four well-known phases (G1, S, G2, and M), for which markers had previously been defined [39]. We considered these markers in this study, and likewise divided phases G1 and S into two early and late subphases as those phases show measurable changes between their beginning and end (Table 2). As for the circadian clock, we identified four different phases corresponding to activity peaks of specific core clock entities (that we subsequently name RORG, SIRT1, BMAL1-CLOCK and PER12-CRY12) [24], that we considered as markers for these phases (Table 3).

**Figure 2:**
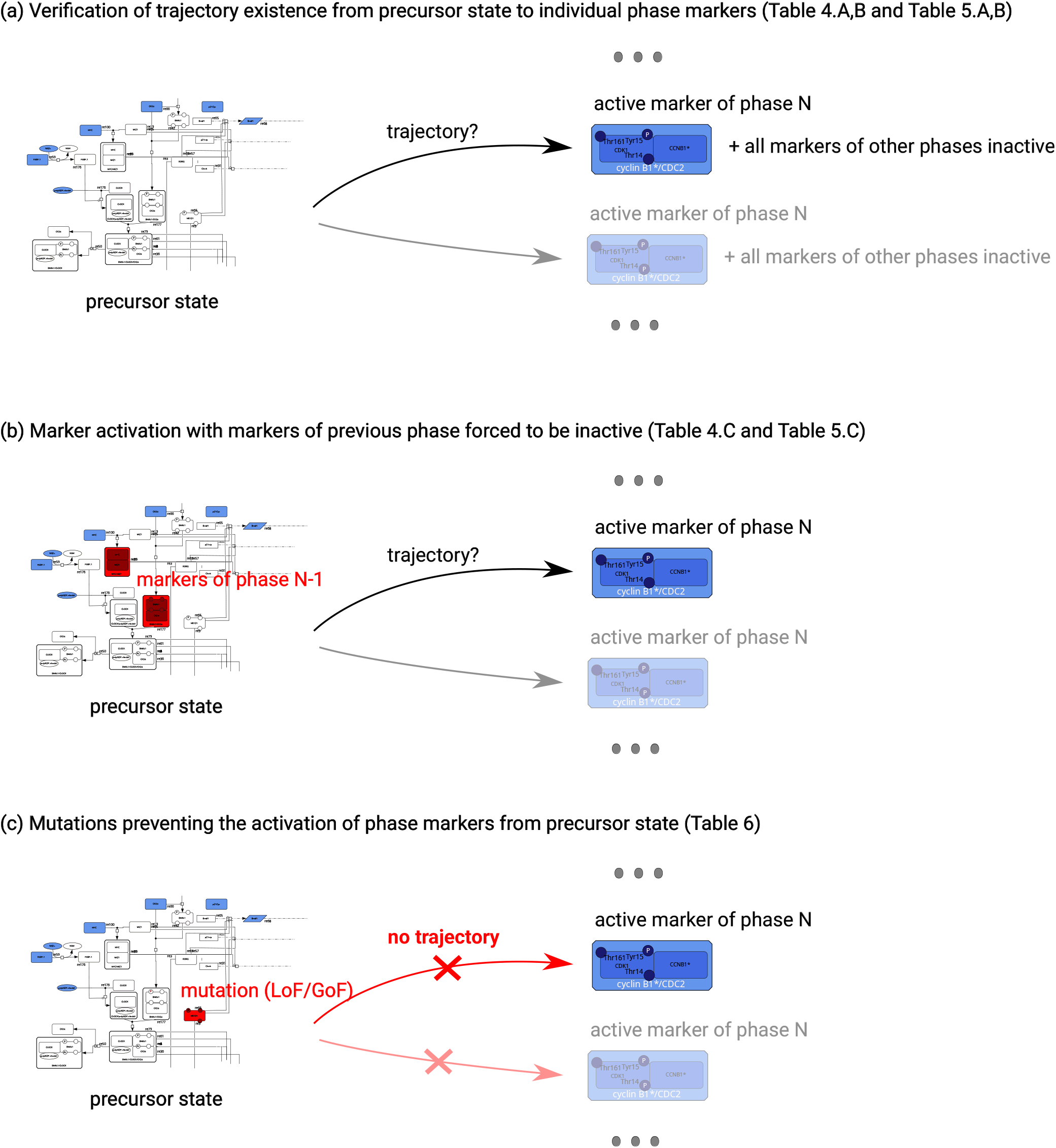
Illustration of the qualitative validations and predictions performed on the dynamical model. The precursor state is composed of the least number of entities to be active (represented in blue) so that the different processes can be activated in the future. The qualitative analyses focus on checking (a,b) or controlling (c) the possibility to activate each marker individually from this precursor state. In (b), the markers of previous phase are disabled, i.e., they can never be activated. In (c), mutations either disables an entity (LoF), or forces its activity (GoF).

**Figure 3:**
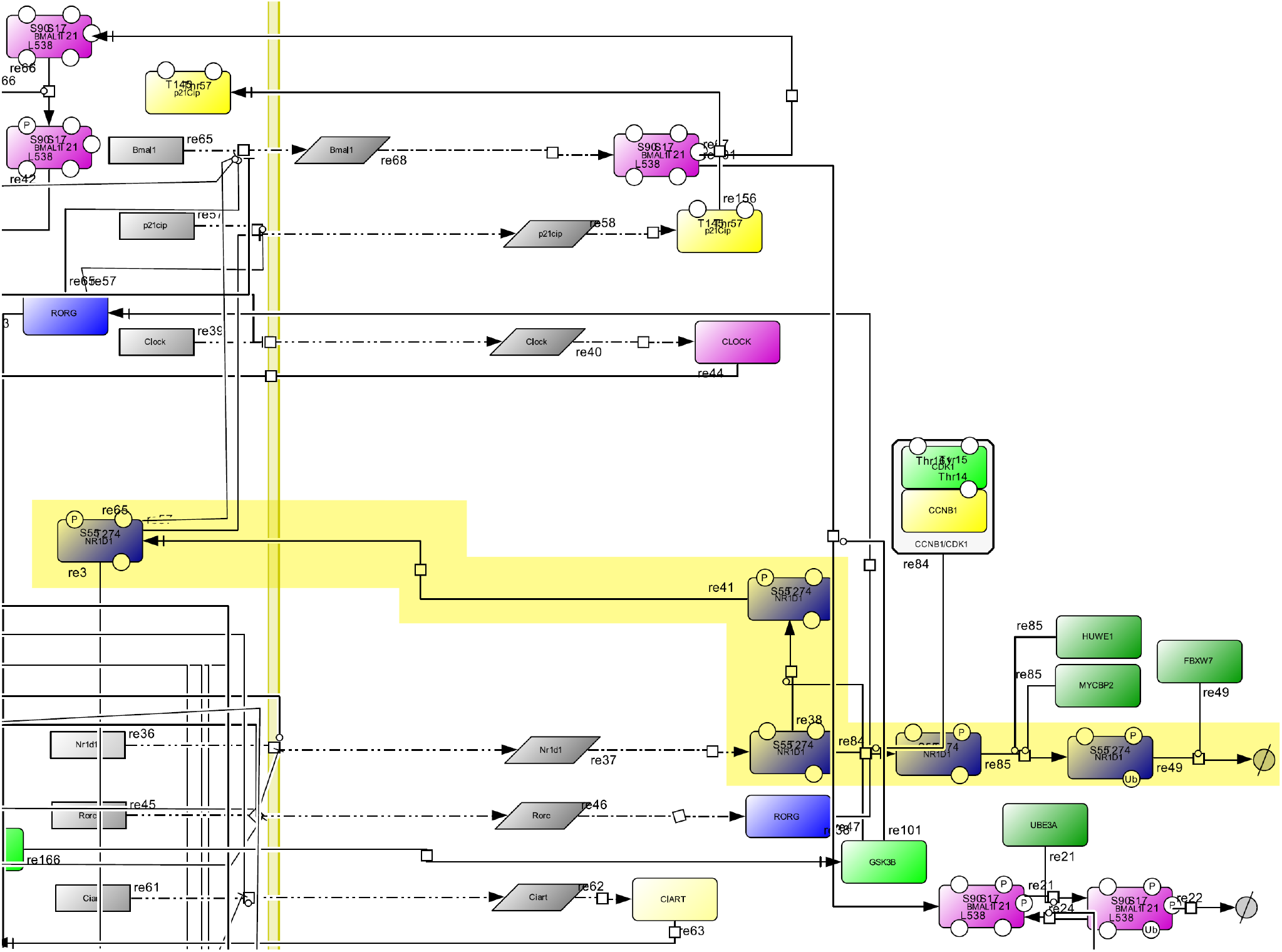
Detailed view of the NR1D1 subnetwork, visualized in CellDesigner. The different forms of NR1D1 are highlighted in yellow. The subnetwork is represented using the CellDesigner format [15], which is compatible with the SBGN Process Description language [40].

**Table 2:** Markers of the phases of the cell cycle. The markers for each phase of the cell cycle are given under a textual form. Text in parantheses gives the post-translational modifications of a given marker or marker’s subunit. For example, “P@Thr172” represents a phosphorylation at threonine 172. A same protein may have more than one post-translational modifications, separated in the text by the “—” character.

| Phase | Marker | Compartment |
| --- | --- | --- |
| early G1 | cyclin D1:CDK6(P@Thr):CDKN1B | nucleoplasm |
|  | pRB:E2F1:DP1:SWI:SNF:HDAC1 | nucleoplasm |
|  | SWI:SNF:HDAC1:SUV39H1:pRB:E2F1:DP1 | nucleoplasm |
|  | cyclin D1:CDK4(P@Thr172):CDKN1B | nucleoplasm |
|  | pRB:E2F1:DP1:SWI:SNF:HDAC1:SUV39H1:HP1gamma | nucleoplasm |
| late G1 | pRB(P):E2F1:DP1:SWI:SNF:HDAC1 | nucleoplasm |
|  | pRB(P—P):E2F1:DP1:SWI:SNF | nucleoplasm |
| early S | cyclin E1:CDK2(P@Thr160) | nucleoplasm |
| late S | cyclin A2:CDK2(P@Thr160) | nucleoplasm |
| G2 | CCNB1:CDK1(P@Tyr15) | cytoplasm |
|  | CCNB1:CDK1(P@Tyr15—P@Thr14) | cytoplasm |
|  | CCNB1:CDK1 | cytoplasm |
|  | CCNB1:CDK1(P@Thr14) | cytoplasm |
| M | CDC25(P) | nucleoplasm |
|  | WEE1(P@Ser53) | nucleoplasm |
|  | CCNB1:CDK1(P@Thr161) | nucleoplasm |
|  | CDC20(P) | nucleoplasm |

**Table 3:**
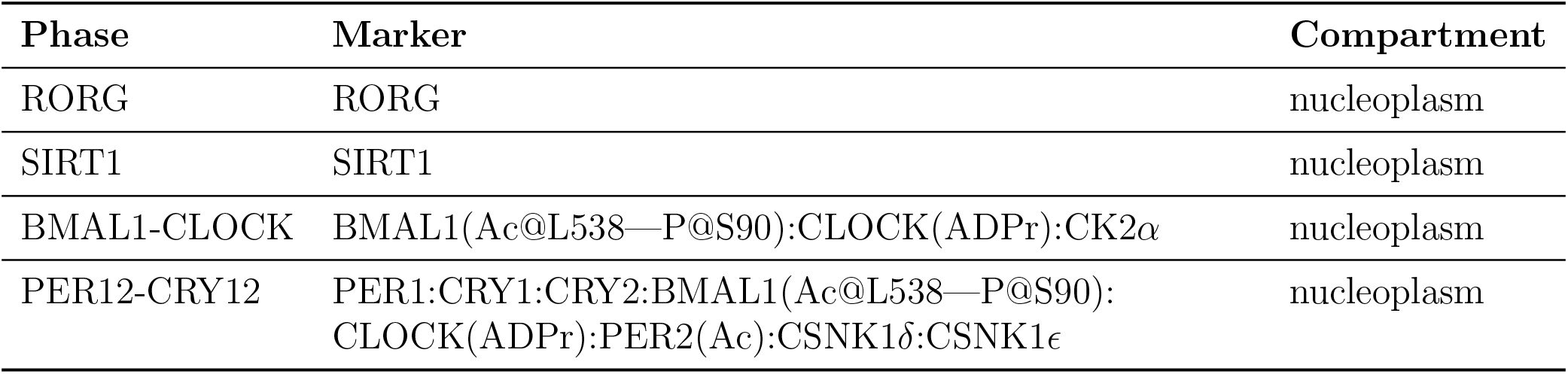
Markers of the phases of the circadian clock. The markers for each phase of the cell cycle are given under a textual form. Text in parentheses gives the post-translational modifications of a given marker or marker’s subunit. For example, “Ac@L538” represents an acetylation at lysine 538. One single protein may have more than one post-translational modifications, separated in the text by the “—” character.

#### 2.3.1 Construction of the dynamical model

We built the dynamical model from the merged map using a qualitative dynamical interpretation, introduced in [39]. In this interpretation, each entity and process of the map is considered as a binary variable whose state (inactive/active, or 0/1) can evolve in time, subject to the state of all entities and processes of the map. For instance, a process can switch from inactive to active whenever all its reactants are active and its modulation logic is satisfied (catalyst active, or inhibitor inactive, for instance). Different rules of state changes exist [39] and may impact substantially the predictions. The interpretation we opted for in this study is general: it does not add constraints between the joined activities of entities modelling different stages of molecular activities. Whereas enabling many behaviors, it provides guarantees with respect to stochastic interpretations of the map, without having to specify kinetics (see Materials and Methods and [13]). The resulting dynamical model is expressed as automata networks, or equivalently 1-safe Petri nets, from which one can make simulations of the evolution of the activity of entities and processes of the map, and make exhaustive analysis of possible trajectories. The model obtained from the merged map resulted in 609 automata and 1220 rules for state changes.

#### 2.3.2 Choice of the initial state

As illustrated by Fig. 2, the validation of the model boils down to verifying the existence of trajectories leading to active markers from a unique initial state of the model, i.e. a set of entities defined being present at the initial time step. In the following, we refer to this unique initial state as the *precursor state*, which has been computed automatically from the merged map (see Materials and Methods for more details).

The definition of the precursor state is critical in such analyses as it has a preponderant effect on reachability results. As the exact coupling of the two cycles has still not been elucidated, it is not possible to define an initial state that would represent an actual state of the coupling that could be observed experimentally. Hence, we built the precursor state mostly grounded on dynamical considerations. The only constraint based on *a priori* knowledge we considered for the construction was the inclusion in the precursor state of the complex formed of the three proteins pRB (unphosphorylated), DP1 and E2F1 rather than of these proteins in their free forms. Indeed, this complex plays a major role in the beginning of the cell cycle: E2F1 becomes active and triggers the transition from G1 to S when it is freed from the complex it forms with pRB subsequently to the phosphorylation of the latter [10]. The rest of the precursor state was built in such a way that it would have a minimal impact on the reachability results. To this end, we chose the initial set among all sets of entities that included the aforementioned complex and that allowed activating successively all processes of the model when present at the initial time step, without considering the effects of modulators. Considering such a constraint for the precursor state guarantees that any absence of reachability is due to internal effects of modulators of the model (i.e., its structure) rather than to the precurosr state itself. Many sets of entities satisfied this second constraint. Hence, we chose a set of entities that both maximized the number of proteins in their most unmodified and free forms and that had the least cardinality. There were 24 such sets; we chose the one containing the entities p53, E2F4 and DP2 (in their most simple forms) located in the cytoplasm and the NAD+ entity located in the nucleoplasm (see Supplementary Information section 2 for a desciption of all alternatives).

#### 2.3.3 Progression of the cell cycle

We checked whether our dynamical model exhibited a correct succession of the phases of the cell cycle. First, we checked whether each phase can be reached from the precursor state. Results are given in Table 4 A. Since each phase could be reached, we subsequently checked whether each phase could be reached independently of each other (i.e., a state with at least one marker of the phase to be reached being present and all markers of the other phases being absent). Again, all phases can be reached (Table 4 B). Finally, to test for a correct succession of phases, we checked whether each phase could be reached when the phase preceding it was disabled. Results are given in Table 4. Early G1 phase can still be reached when M phase is disabled. This could be explained by the choice of the precursor state: because the original cell cycle map [10] was designed as a succession of processes occurring from early G1 phase to M phase, we hypothesize that the precursor state we computed may be close to the states of the cycle corresponding to early G1 phase, and hence that disabling M phase would have no impact on the reachability of early G1. The G2 phase can also still be reached when disabling late S phase. This result is similar to the one we found earlier [39], and can be explained by the fact that the original cell cycle map does not describe any influence of the complex cyclin A2:CDK2, active at the end of S phase, on the activation of the complex cyclin B1:CDK1 (marker of G2). Although such an influence has strong evidence in the literature and could have an important role in the succession late S to G2, its mechanism is still unknown, and hence was not added to the model. All other phases but these two (early G1 and G2) could not be reached when disabling their preceding phase, suggesting that the merged model could reproduce their succession correctly.

**Table 4:**
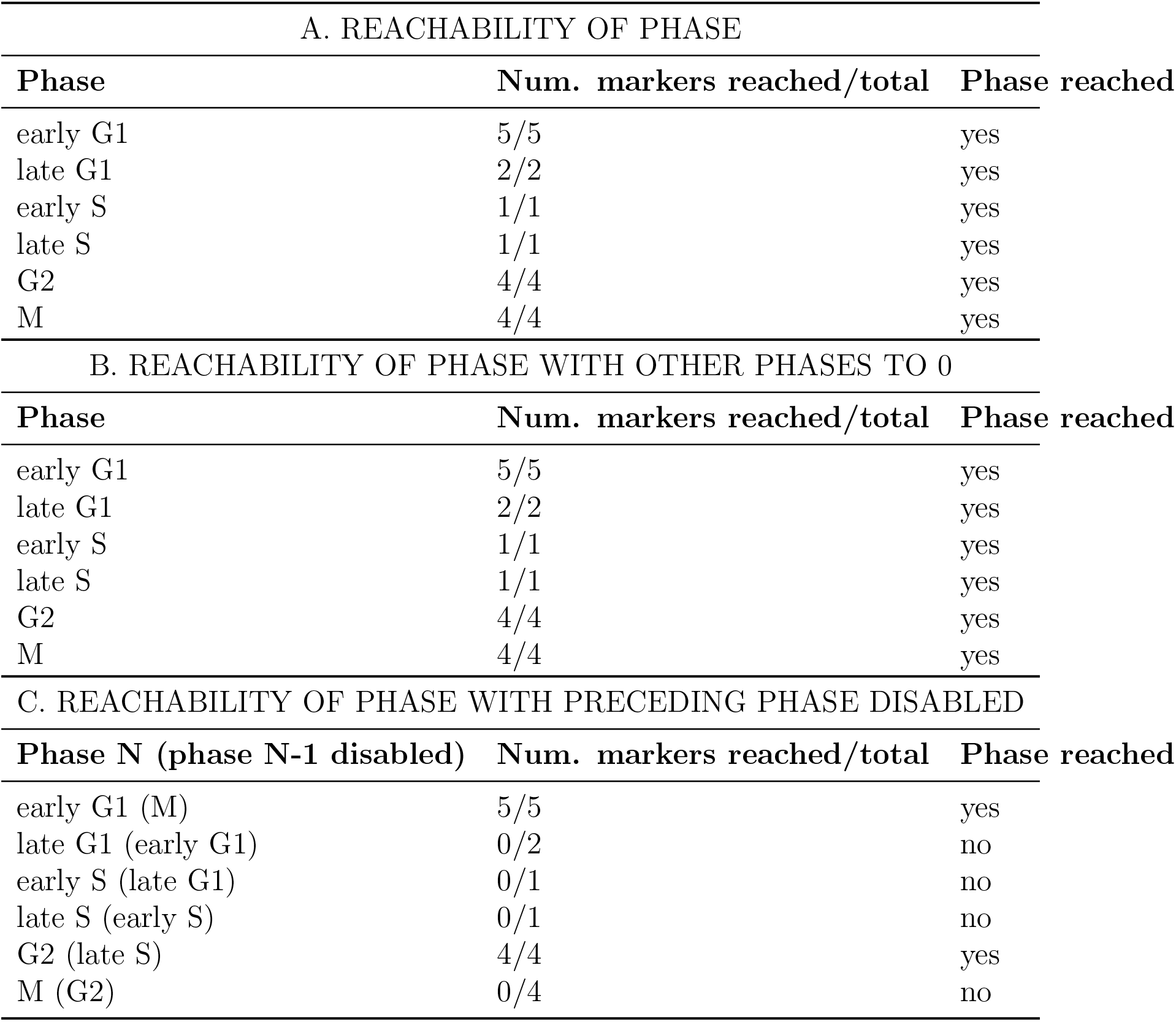
Analysis of the progression of the cell cycle in the dynamical model. We tested whether each phase of the cell cycle could be reached from the precursor state in the model, under different conditions. A phase is reached if at least one of its marker is reached, and is not reached otherwise. A marker is reached if there is a reachable state where the marker is present. A. Phase reached from the precursor state with no other conditions. B. Phase reached from the precursor state while all markers of the other phases are absent. C. Phase N reached from the precursor state, with phase N-1 disabled.

Overall, the results obtained here for the model built from the merged map are the same as those obtained with a model built from the original map of the cell cycle alone [39], showing that the addition in the model of the processes of the circadian clock does not hamper the correct reproduction of the progression of the cell cycle.

#### 2.3.4 Progression of the circadian clock

We conducted the same analysis to check whether our dynamical model exhibited a correct succession of the phases of the circadian clock. We used the same precursor state and the four phases described previously (RORG, SIRT1, BMAL1-CLOCK, and PER12-CRY12). Results are given in Table 5. All phases could be reached from the precursor state (Table 5 A), and all phases could also be reached independantly of all other phases (Table 5 B). Finally, we checked whether each phase could be reached when its preceding phase was disabled. We found that all phases could still be reached except for PER12-CRY12 (Table 5 C). We could expect that RORG would still be reachable: analogously to the cell cycle map, the circadian clock map was designed as a succession of phases from RORG to PER12-CRY12, and we hypothesize that the precursor state is close to the states of the circadian clock when in phase RORG. Hence disabling PER12-CRY12 would have no effect on the reachability of RORG. The fact that SIRT1 could still be reached when disabling RORG can be explained by the absence in the map of an influence of RORG on the activation of SIRT1, which only requires the presence of NAD+. This might suggest we are lacking knowledge regarding the succession from RORG to SIRT1, and more generally on how the activation of SIRT1 is linked to the other processes of the circadian clock. As for the result of reachability for succession SIRT1 to BMAL1-CLOCK, it can be explained by the processes SIRT1 modulates. Indeed, SIRT1 stimulates both the degradation of BMAL1 and PER2, limiting their activity during the phase of SIRT1 and, acting as an inhibitor of the circadian clock. Thus, it could be expected that disabling SIRT1 would not hamper the progression of the circadian clock and the reachability of phase BMAL1-CLOCK.

**Table 5:** Analysis of the progression of the circadian clock in the dynamical model. We tested whether each phase of the circadian clock could be reached from the precursor state in the model, under different conditions. A phase is reached if at least one of its marker is reached, and is not reached otherwise. A marker is reached if there is a reachable state where the marker is present. A. Phase reached from the precursor state with no other conditions. B. Phase reached from the precursor state while all markers of the other phases are absent. C. Phase N reached from the precursor state, with phase N-1 disabled.

| A. REACHABILITY OF PHASE |  |  |
| --- | --- | --- |
| Phase | Num. markers reached/total | Phase reached |
| RORG | 1/1 | yes |
| SIRT1 | 1/1 | yes |
| BMAL1-CLOCK | 1/1 | yes |
| PER12-CRY12 | 1/1 | yes |
| B. REACHABILITY OF PHASE WITH OTHER PHASES SET TO 0 |  |  |
| Phase | Num. markers reached/total | Phase reached |
| RORG | 1/1 | yes |
| SIRT1 | 1/1 | yes |
| BMAL1-CLOCK | 1/1 | yes |
| PER12-CRY12 | 1/1 | yes |
| C. REACHABILITY OF PHASE WITH PRECEDING PHASE DISABLED |  |  |
| Phase N (phase N-1 disabled) | Num. markers reached/total | Phase reached |
| SIRT1 (RORG1) | 1/1 | yes |
| BMAL1-CLOCK (SIRT1) | 1/1 | yes |
| PER12-CRY12 (BMAL1-CLOCK) | 0/1 | no |
| RORG (PER12-CRY12) | 1/1 | yes |

#### 2.3.5 Control of checkpoints

How mechanisms described in the circadian clock map may affect the progression of the cell cycle, and vice-versa? This question is central for understanding the potential intertwined control of the two cycles. Using qualitative dynamics, we addressed this question by looking for processes controlling the reachability of the different cycle checkpoints, specified by markers. Due to the size of the model, we used methods based on logical deduction to avoid a naive screening of all possible candidates, and relied on formal approximations of dynamics, which result in a correct but non-exhaustive list of controls (see Materials and Methods).

Our analysis identified sets of simultaneous mutations, being either Loss-of-Function (LoF) or Gain-of-Function (GoF), which ensure that one marker of a checkpoint is no longer reachable from the precursor state. We report here the entities from the circadian clock map that are involved in the sets of mutations interfering with the checkpoints of the cell cycle, and similarly, the entities of cell cycle map interfering with the checkpoints of the circadian clock.

##### Control of cell cycle checkpoints by entities of the circadian clock map

Starting from the early G1 state, Table 6 lists the mutations of entities of the circadian clock map which can prevent the activation of at least one marker of each phase of the cell cycle. Blockade of the reachability of early G1, late G1 and late S markers upon HSP90 LoF is likely to result from the direct role of the protein in the cell cycle process. HSP90 is a chaperone protein that is required for the maturation and stability of a large number of proteins including entities of the cell cycle machinery such as CDK4 and CDC37 present in the cell cycle map [8]. Further, pharmacological inhibition of HSP90 with 17-Allylamino-Geldanamycin and its derivatives has an antitumor activity in preclinical models and cancer patients [16, 22]. LoF preventing reachability of G2 and M markers includes mutations impairing directly or indirectly (CK2*α*, PARP1, MYC and MIZ1) CLOCK/BMAL1 function which is the core transcription factor complex driving the circadian clock mechanism and the expression of the G2/M kinase WEE1 [29]. LoF of CLOCK/BMAL1 function also leads to increased expression of the CDK4/6 inhibitor p21 as the NR1D1 transciptional repressor is no longer produced [21]. The impact of WEE1 LoF on G2 marker reachability lies in its role as the kinase that phosphorylates CDK1 at Tyr15 during G2. Intriguingly, this analysis revealed that NR1D1 GoF participates in preventing activation of one of the G2 and M markers. According to the analysis of the merged map, the activation of NR1D1 can prevent the activation of the complex PER:CRY:CSNK1:CLOCK:BMAL1, which is necessary for the activation of the CCNB1:CDK1 phase markers of G2 and M. This is in line with data showing that experimental overexpression of NR1D1 in vivo inhibits the expression of BMAL1 [23]. Expectedly, LoF for CCNB1:CDK1 which is the key driver of the G2/M transition blocked the reachability of phase M markers.

**Table 6:**
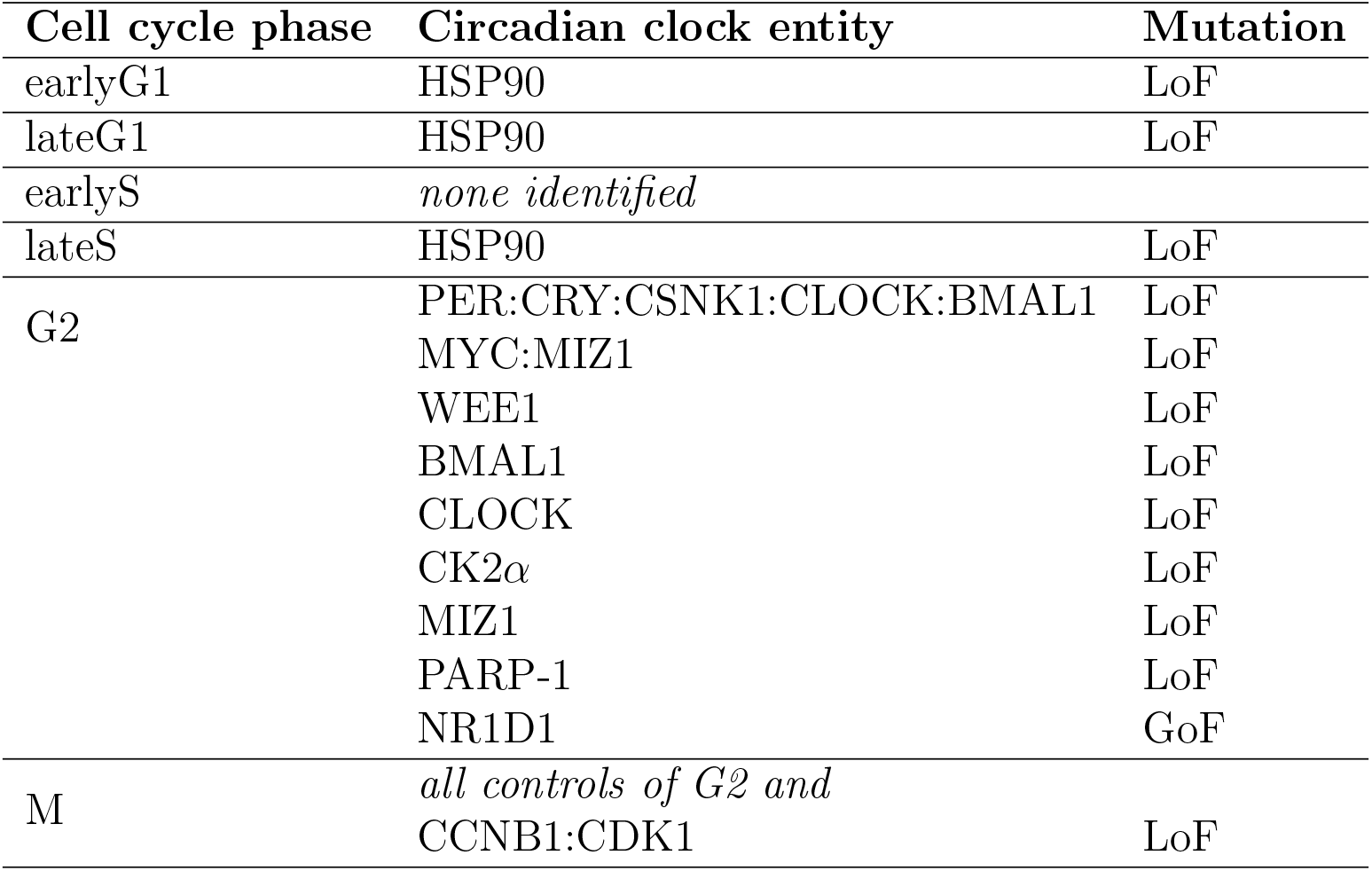
Identified mutations of entities of the circadian clock controlling the reachability of at least one phase marker of a cell cycle phase in the dynamical model.

| Cell cycle phase | Circadian clock entity | Mutation |
| --- | --- | --- |
| earlyG1 | HSP90 | LoF |
| lateG1 | HSP90 | LoF |
| earlyS | *none identified* |  |
| lateS | HSP90 | LoF |
| G2 | PER:CRY:CSNK1:CLOCK:BMAL1 | LoF |
|  | MYC:MIZ1 | LoF |
|  | WEE1 | LoF |
|  | BMAL1 | LoF |
|  | CLOCK | LoF |
| | CK2α | LoF |
|  | MIZ1 | LoF |
|  | PARP-1 | LoF |
|  | NR1D1 | GoF |
| M | *all controls of G2 and* |  |
|  | CCNB1:CDK1 | LoF |

##### Control of circadian clock checkpoints by entities of the cell cycle map

The qualitative analysis identified no mutations of cell cycle entities involved in the blocking of circadian clock markers. However, a causal and exhaustive analysis of the trajectories leading to markers activation revealed that several trajectories leading to the activation of RORG, BMAL1-CLOCK, and PER-CRY from the precursor state rely on the activity of p53. This link is supported by an experimentally documented bidirectional crosstalk between the circadian clock and the p53 pathway that modulates the stability of the PER2 and p53 proteins [27, 48].

## 3 Discussion

### 3.1 Detailed map of the circadian clock and its merging with the mammalian cell cycle map

We built a new map of the circadian clock that encompasses all known key regulatory mechanisms governing the core clock, as well as key entities that also play a major role in the mammalian cell cycle. To our knowledge, this map is the most detailed and up-to-date map made publicly available as compared to the currently existing maps from Reactome (https://reactome.org), PANTHER (http://www.pantherdb.org), and KEGG (https://www.genome.jp/kegg) databases. The circadian clock map from Reactome map includes a moderately detailed core clock and numerous clock outputs while the ones from Panther and KEGG describe a minimalist core clock with 6 and 13 nodes respectively. Compared to these two existing resources, the present map focuses on the core clock components, includes important synchronisation input pathways and nodes present in other critical cellular processes (cell cycle, metabolism), and incorporates a detailed and updated information related to numerous posttranslational modifications (phosphorylation, acetylation, poly-ADP-ribosylation, and ubiquitination). The map was designed using CellDesigner [15] and is compatible with the SBGN PD standard [40]. As such, it may be easily shared, edited, and updated. Its referencing and integration in map databases, such as the Atlas of Cancer Signalling Network [26], is under consideration.

We then merged this map to the mammalian cell cycle map, which describes the mechanisms governing the regulation of the cycle by key regulators such as the E2F family and the pRB protein. The merging required making changes to the two maps to make them compatible, namely renaming, modifying, deleting or adding entities in the two maps. It was performed programmatically on the conceptual models underlying the two maps using the *csbgnpy* Python library, which allowed making all changes explicit without altering the source files of the input maps.

### 3.2 Validation of the map and its coupling by qualitative dynamical analysis

We used qualitative approaches to ensure that the merged map provides a consistent description of the molecular processes sufficient to reproduce expected basic dynamical behavior of both cycles (the mammalian cell cycle and the circadian clock) and of their coupling. For this, we built a formal dynamical model from the merged map, relying on a specific dynamical interpretation of SBGN PD maps and encoded under the form of automata networks (framework that is related to Petri nets) [39]. The interpretation we used precludes the need for kinetic parameters, and avoids fixing arbitrary logics for modulations, so as not to arbitrarily exclude specific dynamical behaviors. We then analyzed the dynamics generated by our model using the *Pint* software tool. Specifically, we verified the correct progression of both cycles by verifying the existence of trajectories leading to phase markers of each cycle, and predicted mutations to control these trajectories. Our model could reproduce a correct progression of the cell cycle for all but two phases (M to early G1 and late S to G2), which could be explained by the way the map and the precursor state are built (for M to early G1) and a lack of knowledge on the precise mechanisms leading to the activation of the complex cyclin B1:CDK1 by the complex cyclin A2:CDK2 at the end of phase S (for late S to G2). Additionnally, our model could reproduce a correct progression from BMAL1-CLOCK to PER12-CRY12 phases, a succession which underlies the core negative transcriptional/postranslational feedback loop of the clock mechanism. However, it failed to reproduce a correct progression for the rest of the phases (PER12-CRY12 to RORG, RORG to SIRT1 and SIRT1 to BMAL1-CLOCK), due to the way the map and precursor state are built (PER12-CRY12 to RORG), the absence of an influence of RORG on SIRT1 (RORG to SIRT1) and to the overall inhibitory influence of SIRT1 on the circadian clock (SIRT1 to BMAL1-CLOCK). These latter two results might reveal a lack of knowledge surrounding marker SIRT1. If its effects on the circadian clock are well known (inhibition by inhibiting BMAL1 and PER2), the processes that come into play for its activation at phase RORG are still not clearly understood, and were consequently not taken into account in our model.

### 3.3 Insights in the circadian clock and cell cycle interactions

Using the merged map, we analysed how entities and processes brought by the circadian clock map can control the progression of the cell cycle, and vice versa. The analysis relied on the identification of mutations which prevent the activation of a marker of a phase. We recovered expected regulatory mechanisms of the circadian clock to the cell cycle, and identified NR1D1 as a potential inhibitor for the progression towards the G2 phase. This prediction corroborates observations made in the literature showing that two agonists of NR1D1 are lethal to cancer cells and oncogene-induced senescent cells [43, 46], and calls for further investigation.

As for the analysis of the influence of the cell cycle map on the circadian clock map progression, it resulted in very few predictions. This reflects the fact that there are few known mechanistic effects from the cell cycle on the circadian clock.

### 3.4 Future directions

Further analysis of the dynamics of both cycles and their coupling would necessitate studying some of their temporal properties, such as their period. These properties cannot be easily analyzed using qualitative approaches, because they overlook quantitative time. Nevertheless, inhibition processes, which can regulate the duration of phase transitions, may still be captured by causal analysis by identifying entities involved in pathways of phase progressions, e.g. with cut set analysis [37], together with processes de-activating these entities. Furthermore, the present analysis is restricted to the study of trajectories from a *precursor state* from which all the entities of the map may be activated. Being able to delineate network states corresponding to different cycle phases would allow providing further insight in key entities and processes regulating the circadian clock and cell cycle progression, and their synchronization.

In future work, we plan to study alternative dynamical interpretations of SBGN PD maps, such as CasQ [1]. This interpretation results in a model expressed in the classical Boolean network framework, where tools such as Most Permissive Boolean Networks [38] can address the dynamical analysis of networks of the scale of the merged map. It would notably help providing a finer and complete analysis of trajectories, long-term behaviors (attractors), and mutations.

## 4 Materials and Methods

### 4.1 SBGN PD standard for maps

The Systems Biology Graphical Notation (SBGN) [28] is a standard for the graphical representation of molecular networks. It includes three orthogonal languages: the Process Description (PD) [40], used to represent reaction networks; the Activity Flow [30], used to represent influence graphs; and the Entity Relationship [42], used to represent sets of rules and relationships. We focus hereafter on the SBGN PD language, as we are interested in representing and modelling reaction networks.

SBGN PD allows representing the precise biomolecular mechanisms underlying biological processes. As its name suggests, the central concept it permits representing is the process, that transforms pools of entities into other pools of entities, and that may be modulated by yet other pools of entities. PD defines a number of glyphs (nodes and arcs), each having a specific shape and representing a different concept. The main glyphs of the PD language are the following:

- *Entity Pool Node (EPN)*: represents a pool of entities, that is a set of identical molecules. There are 13 types of EPNs, among which the macromolecule, simple chemical, complex and nucleic acid feature.
- *Compartment Node*: represents a compartment, where entity pools are localized.
- *Process Node*: represents a process that transforms one or more entity pools into other entity pools. The reactants and products of the process are linked to it using consumption arcs and production arcs, respectively. There are six types of processes: the generic process, association, dissociation, phenotype, omitted process and uncertain process.
- *Modulation arc*: represents the modulation of a process by an entity pool. There are five types of modulation arcs: the generic modulation, stimulation, inhibition, necessary stimulation, and catalysis.
- *Logical Operator Node*: allows building complex logical function that model complex modulations (e.g. cooperation may be modelled with and And operator). There are three types of logical operator nodes: the and operator, or operator, and not operator.

An actual drawing in SBGN of a network is called an SBGN map. SBGN maps may be stored and exchanged under the SBGN-ML format [5]. Multiple software allow drawing such maps [44], one of which is CellDesigner [15]. This particular software focuses on the edition of SBGN PD maps and has been used to draw large comprehensive maps [26, 31, 34, 33, 10]. It only partially supports the SBGN-ML format and mostly relies on its own format to import and export maps. This format is an extension of the SBML format, and can be converted to SBGN-ML using the *cd2sbgnml* software [2].

### 4.2 Construction of the circadian clock map

The biological knowledge providing a comprehensive understanding of the molecular circuitry governing the mammalian circadian clock available on December 2018 was used to design a prototypical peripheral circadian clock map in CellDesigner 4.4.2 (http://www.celldesigner.org) using the SGBN standard (https://www.sbgn.org). This map did not intend to exhaustively describe all the known species and interactions but instead incorporate essential nodes and edges of the core clock mechanism as well as critical inputs and outputs connecting the core clock and the cell cycle machinery. Accordingly, paralogs were not included in most instances and specific accessory loops (DBP, DEC1) were also omitted in this version. As eukaryotic circadian clocks are based on interwoven transcriptional/translational feedback loops, the circadian clock map incorporates genes, mRNA and protein products with their associated posttranslational modifications (phosphorylation, ubiquitination, acetylation). However, when transcriptional regulation was not essential or not documented, only proteins were considered for some modules. It also incorporates input pathways known to be important for the entrainment of peripheral clocks such as glucocorticoid signaling and temperature. Genetic interactions not supported by mechanistic evidence have not been included. This map compiles only experimentally validated results obtained mostly in the mouse animal model. The map is enriched with 297 external links to the Entrez, UniprotKB, ChEBI and PUBMED databases.

### 4.3 Modification and merging of the cell cycle and circadian clock maps

In order to modify and merge the cell cycle and circadian clock maps, we developed *csbgnpy*, a Python library that eases the manipulation of SBGN PD maps, at the conceptual level. With *csbgnpy*, maps can be modified programmatically without alteration of the input files, and the applied modifications can be easily tracked. *csbgnpy* extracts all the biological and biochemical concepts represented by a map (e.g. entity pools, processes, modulations) and stores them in memory under the form of Python objects that can be manipulated using simple Python code. The operations facilitated by *csbgnpy* include the creation of new concepts and their addition to a map; the search for concepts in a map; the deletion or modification of concepts of a map; and the construction of new maps by merging or intersecting two or more maps. Concepts in *csbgnpy* may also be manipulated using a new textual representation called sbgntxt, that eases their construction or identification. *csbgnpy* can read maps from different file formats such as SBGN-ML, CellDesigner and BioPAX (using third-party libraries or converters, in particular *cd2sbgnml* [2] for the CellDesigner format) and write maps to the SBGN-ML format (with no layout). *csbgnpy* is freely available at https://github.com/Adrienrougny/csbgnpy, and can be modified and redistributed under the terms of the GPLv3 license.

The modification and merging of the two maps, as well as the export of the resulting map to an SBGN-ML file, were performed programmatically using *csbgnpy*, directly from the input CellDesigner files. All changes made to both maps are described in details in Supplementary Information section 1.

### 4.4 Dynamical model

#### 4.4.1 Construction of the dynamical model

We built a qualitative dynamical model from the merged map using the *sbgnpd2an* program. This program takes as input an SBGN PD map stored under the SBGN-ML format and outputs a dynamical model stored under the *Pint* format [35] using the semantics and the methods described in [39]. This method allows building a qualitative dynamical model from an SBGN PD map, and encoding it under the form of an asynchronous automata network (AN). We used the general semantics, introduced in [39], to build our model. We briefly describe this semantics and the encoding under the form of an AN hereafter.

The general semantics is a qualitative Boolean semantics that extends the one of BIOCHAM [9]. Under this semantics, (i) each EPN may be either present or absent; (ii) each process may be either occurring or non-occurring; (iii) each modulation may be either active or inactive. Moreover, (iv) a modulation is active if its source is satisfied, and inactive otherwise. If the source of the modulation is a unique EPN, it is satisfied if and only if this EPN is present; whereas if the source is a logical function structuring some EPNs, it is satisfied if and only if the logical function is satisfied, where an EPN that is present counts for true and an EPN that is absent for false; (v) a process may become occurring if and only if all its reactants are present, all the necessary stimulations targeting it are active, and at least one stimulation targeting it is active or one inhibition targeting it is inactive; (vi) a process may become non-occurring if and only if all its products are present; (vii) an EPN may become present if and only if it is the product of an occurring process; (viii) and finally, an EPN may become absent if and only if it is the reactant of an occurring process and all the products of this process are present. This semantics is permissive: the constraint to be satisfied for a process to become occurring is the weakest possible when still considering the usual meaning of stimulations and inhibitions, and a reactant of a process may never be consumed even if the products of the process have been produced. A model built using this semantics may be encoded under the form of an AN as follows: (a) each EPN is encoded by one automaton that has two local states ‘1’ and ‘0’, encoding the presence and the absence of the EPN, respectively; (b) each process is also encoded by one automaton with local states ‘1’ and ‘0’, encoding this time the occurrence and non-occurrence of the process, respectively; (c) and transitions between local states of automata are encoded and conditioned straightforwardly from elements (iv-viii) of the semantics. For example, an automaton encoding an EPN will have one transition from its state ‘0’ to ‘1’ for each process that produces it, and this transition will be conditioned by the automata modelling the producing process being in its local state ‘1’, following (vii). The update scheme considered is the asynchronous one, where only one transition may be fired at each time step (in terms of the model, only one process or EPN may change its state at each time step), making the entailed dynamics non-deterministic. Figure 4 shows an AN-encoded model built from an example SBGN PD map under the general semantics.

**Figure 4:**
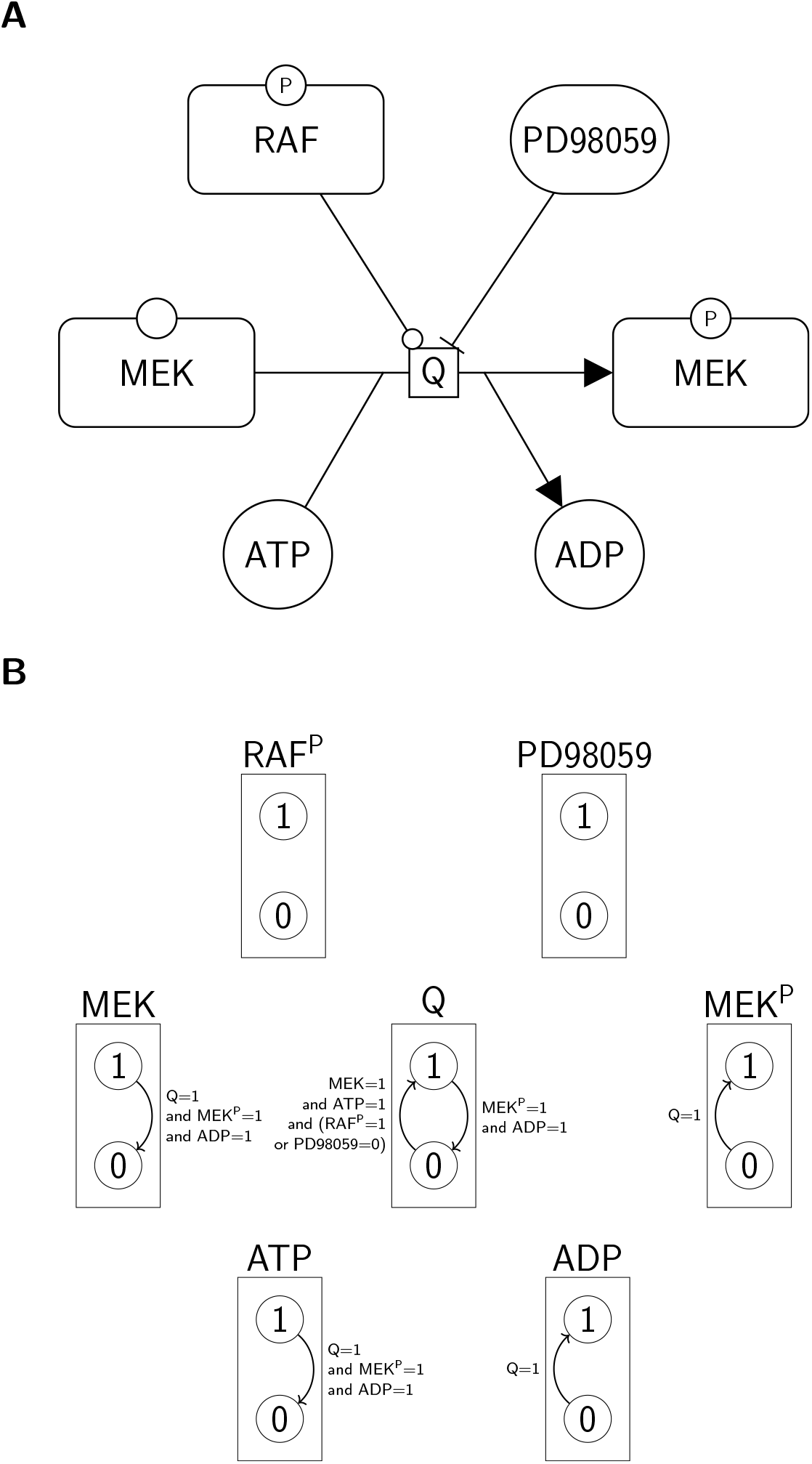
An AN-encoded model built from an example SBGN PD map under the general semantics. A. An example SBGN PD map that represents the catalysis of the phosphorylation of MEK by p-RAF and its inhibition by PD98059. B. The AN that encodes the model built from the SBGN PD map under the general semantics. Each rectangle box represents an automaton, and the circles inside the automata represent their local states. Arrows represent transitions between local states, and are conditioned by the states of some automata of the network.

#### 4.4.2 Model analysis

##### Construction of the precursor state

The precursor state used for our analyses was computed automatically using Answer Set Programming (ASP) [17] and the software *clingo* [18]. Our goal was to generate a minimal subset of entities (forming the precursor state) that would allow to produce all entities of the map not considering modulations and that would maximize the total number of entities in their free and unmodified states. For this, we built an ASP program encoding the reaction graph underlying the merged map (with all modulations removed) and the following rules and constraints. We say that an entities is *covered* if it can be produced from the precursor state:

- *Rule 1* : an entity might be in the precursor state or not (generative rule);
- *Rule 2* : if an entity is in the precursor state, then it is covered;
- *Rule 3* : if an entity has a process producing it and consuming an empty set (source), then it is covered;
- *Rule 4* : if an entity is produced by a process whose reactants are covered, then it is covered;
- *Constraint 1* : all entities must be covered;
- *Constraint 2* : the precursor state must contain the complex formed of the three proteins pRB (unphosphorylated), DP1 and E2F1;
- *Optimization 1* : minimize the total number of entities;
- *Optimization 2* : minimize the total number of state variables set to a value;
- *Optimization 3* : minimize the total number of complexes.

The built ASP program was then solved using *clingo*, and the precursor state was selected among the 24 possible solutions.

##### Reachability analysis

Given a dynamical model, the reachability analysis consists in checking if there exists a trajectory from a given initial state of the model to another state where a given set of markers are present. For asynchronous automata networks (similarly to asynchronous Boolean networks), the search of such trajectories has a high computational complexity. We rely on the software *Pint* [35], which implements a method to reduce the model while preserving the desired reachability property [36]. The dynamically reduced models can then be tractable for exact reachability analysis.

##### Phase succession analysis

The phase succession analysis was performed using software *Pint* on the qualitative model generated from the merged map. Each phase was defined by a number of markers that are supposed to be present in that phase. Hence, a phase was considered reachable if the present state of at least one of its marker was reachable; and phases were disabled by removing all local transitions in-going, out-going, or being conditioned by the present state of each of their markers.

##### Mutations analysis

We used the software *Pint* for the deduction of combinations of LoF/GoF mutations making a reachability property false: after applying the mutation, it is guarantees that no trajectory exist between the specified initial state and target active markers. *Pint* relies on static analysis and logic programming techniques which scale to large networks. The analysis returns a list of combinations of mutations, which may be non exhaustive. In our checkpoint analysis, we report each entity of map A which appears in at least one combination of mutations for blocking the reachability of a marker of map B.

##### Causal analysis of trajectories

We used the software *Pint* to extract entities whose activities are necessary in at least one trajectory between a given initial state and a state containing the given markers. The computation relies on a causal analysis of transitions, and returns sets of entities such that each trajectory relies on a at least one entity of that set.

### 4.5 Data and software availability

The circadian clock map introduced in this article is provided in the *CellDesigner* format in the Supplementary Materials (CircadianClock.xml).

The complete computational analysis, from the merge of the circadian clock and cell cycle maps to the qualitative model validation and mutation predictions, can be repeated using the notebooks available at https://github.com/pauleve/ClockCycle-notebooks and the Docker image *pauleve/clock-cycle:v0*, based on the CoLoMoTo Docker distribution [32] (see https://github.com/pauleve/ClockCycle-notebooks for instructions). It was run on a desktop computer with a 3.3Ghz CPU and 64GB of RAM with a total execution time of 2.5 minutes.

The merge of the circadian clock and cell cycle maps has been performed with the *csbgnpy* Python library in version 0.1, available at https://github.com/Adrienrougny/csbgnpy. The qualitative interpretation of the coupled map resulting in a formal dynamical model expressed in the autonama network framework has been performed with the *sbgn2an* Python library in version 0.1, available at https://github.com/Adrienrougny/sbgn2an and the *pypint* Python library in version 1.6.1, available at https://loicpauleve.name/pint. The qualitative analysis, including model validation and mutation predictions, has been performed using the software *Pint* in version 2019-05-24, available at https://loicpauleve.name/pint.

## 5 Conclusions

In this study, we introduce a new map of the circadian clock, compatible with the SBGN standard. Key features of this map include an unprecedented level of detail, upgrading and merging capabilities and, direct use for dynamic modelling. We programmatically merge this map with the previously published map of the mammalian cell cycle, and automatically build a qualitative dynamical model from the merged map we obtain. We then validate this model by showing it can reproduce the progression through their main phases of both cycles. We subsequently use our model to make predictions on mutations that would influence the coupling of the two cycles, and notably predict that NR1D1 should inhibit the progression of the cell cycle towards phase G2. This result is corroborated by previously reported observations following which pharmacological activation of NR1D1 inhibits tumour cell proliferation, and should be investigated experimentally later. Furthermore, our study relies on generic tools for the merging and the dynamical analysis that are independent of the specific maps introduced, and could be repeated on updated versions of the cell cycle and circadian clock maps, or on maps related to completely different biological processes.

## Supporting information

File CircadianClock.xml

## Supplementary material

The following are available: Supplementary Information section 1: Modification of the maps for the merging, Supplementary Information section 2: Modification of the circadian clock map, File CircadianClock.xml: Map of the circadian clock in the CellDesigner format.

## Authors’ contributions

M.T. and F.D. designed the circadian clock map; A.R. wrote software for SBGN map merging; A.R. and L.P. performed the model analysis; A.R., L.P, F.D. wrote the initial draft of the manuscript. A.R., L.P., M.T., F.D. edited the manuscript.

## Funding

This research was funded by the French Agence Nationale pour la Recherche (ANR): HYCLOCK project (ANR-14-CE09-0011), iCycle project (ANR-16-CE33-0016-01), AlgoReCell project (ANR-16-CE12-0034), “Investments for the Future” LABEX SIGNALIFE (ANR-11-LABX-0028-01), UCAJEDI Investments in the Future project (ANR-15-IDEX-01) and the Canceropôle Provence-Alpes-Côte d’Azur, the French National Cancer Institute (INCa) and the Provence-Alpes-Côte d’Azur Region.

## Conflicts of interest

The authors declare no conflict of interest.

## Abbreviations

The following abbreviations are used in this manuscript:

AN: Automata network
ASP: Answer Set Programming
EPN: Entity Pool Node
GoF: Gain-of-Function
LoF: Loss-of-Function
PD: Process Description
SBGN: Systems Biology Graphical Notation

## Notes

### Competing Interest Statement

The authors have declared no competing interest.

## References

[1] Sara Sadat Aghamiri, Vidisha Singh, Aurélien Naldi, Tomáš Helikar, Sylvain Soliman, and Anna Niarakis. Automated inference of boolean models from molecular interaction maps using CaSQ. Bioinformatics, 36(16):4473–4482, 2020.

[2] Irina Balaur, Ludovic Roy, Alexander Mazein, S Gökberk Karaca, Ugur Dogrusoz, Emmanuel Barillot, and Andrei Zinovyev. cd2sbgnml: bidirectional conversion between celldesigner and sbgn formats. Bioinformatics, 36(8):2620–2622, 2020.

[3] Annabelle Ballesta, Pasquale F Innominato, Robert Dallmann, David A Rand, and Francis A Lévi. Systems chronotherapeutics. Pharmacological reviews, 69(2):161–199, 2017.

[4] Aurélio Balsalobre, Steven A Brown, Lysiane Marcacci, Francois Tronche, Christoph Kellendonk, Holger M Reichardt, Günther Schütz, and Ueli Schibler. Resetting of circadian time in peripheral tissues by glucocorticoid signaling. Science, 289(5488):2344–2347, 2000.

[5] Frank T Bergmann, Tobias Czauderna, Ugur Dogrusoz, Adrien Rougny, Andreas Dräger, Vasundra Touré, Alexander Mazein, Michael L Blinov, and Augustin Luna. Systems biology graphical notation markup language (sbgnml) version 0.3. Journal of integrative bioinformatics, 17(2-3), 2020.

[6] Jonathan Bieler, Rosamaria Cannavo, Kyle Gustafson, Cedric Gobet, David Gatfield, and Felix Naef. Robust synchronization of coupled circadian and cell cycle oscillators in single mammalian cells. Molecular systems biology, 10(7):739, 2014.

[7] Ethan D Buhr, Seung-Hee Yoo, and Joseph S Takahashi. Temperature as a universal resetting cue for mammalian circadian oscillators. science, 330(6002):379–385, 2010.

[8] Francis Burrows, Hong Zhang, and Adeela Kamal. Hsp90 activation and cell cycle regulation. Cell Cycle, 3(12):1530–1536, 2004.

[9] Laurence Calzone, François Fages, and Sylvain Soliman. Biocham: an environment for modeling biological systems and formalizing experimental knowledge. Bioinformatics, 22(14):1805–1807, 2006.

[10] Laurence Calzone, Amélie Gelay, Andrei Zinovyev, François Radvanyi, and Emmanuel Barillot. A comprehensive modular map of molecular interactions in rb/e2f pathway. Molecular systems biology, 4(1):0174, 2008.

[11] Shaon Chakrabarti and Franziska Michor. Circadian clock effects on cellular proliferation: Insights from theory and experiments. Current Opinion in Cell Biology, 67:17–26, 2020.

[12] Rukeia El-Athman and Angela Relógio. Escaping circadian regulation: an emerging hallmark of cancer? Cell systems, 6(3):266–267, 2018.

[13] François Fages and Sylvain Soliman. Abstract interpretation and types for systems biology. Theoretical Computer Science, 403(1):52–70, 2008.

[14] Céline Feillet, Peter Krusche, Filippo Tamanini, Roel C Janssens, Mike J Downey, Patrick Martin, Michèle Teboul, Shoko Saito, Francis A Lévi, Till Bretschneider, et al. Phase locking and multiple oscillating attractors for the coupled mammalian clock and cell cycle. Proceedings of the National Academy of Sciences, 111(27):9828–9833, 2014.

[15] Akira Funahashi, Yukiko Matsuoka, Akiya Jouraku, Mineo Morohashi, Norihiro Kikuchi, and Hiroaki Kitano. Celldesigner 3.5: a versatile modeling tool for biochemical networks. Proceedings of the IEEE, 96(8):1254–1265, 2008.

[16] Rocio Garcia-Carbonero, Amancio Carnero, and Luis Paz-Ares. Inhibition of hsp90 molecular chaperones: moving into the clinic. The Lancet Oncology, 14(9):e358–e369, 2013.

[17] Martin Gebser, Roland Kaminski, Benjamin Kaufmann, and Torsten Schaub. Answer set solving in practice. Synthesis lectures on artificial intelligence and machine learning, 6(3):1–238, 2012.

[18] Martin Gebser, Roland Kaminski, Benjamin Kaufmann, and Torsten Schaub. Clingo= asp+ control: Preliminary report. arXiv preprint arXiv:1405.3694, 2014.

[19] Sigal Gery, Naoki Komatsu, Lilit Baldjyan, Andrew Yu, Danielle Koo, and H Phillip Koeffler. The circadian gene per1 plays an important role in cell growth and dna damage control in human cancer cells. Molecular cell, 22(3):375–382, 2006.

[20] Tetsuya Gotoh, Jae Kyoung Kim, Jingjing Liu, Marian Vila-Caballer, Philip E Stauffer, John J Tyson, and Carla V Finkielstein. Model-driven experimental approach reveals the complex regulatory distribution of p53 by the circadian factor period 2. Proceedings of the National Academy of Sciences, 113(47):13516–13521, 2016.

[21] Aline Gréchez-Cassiau, Béatrice Rayet, Fabienne Guillaumond, Michèle Teboul, and Franck Delaunay. The circadian clock component bmal1 is a critical regulator of p21waf1/cip1 expression and hepatocyte proliferation. Journal of Biological Chemistry, 283(8):4535–4542, 2008.

[22] Wei He and Huixian Hu. Biib021, an hsp90 inhibitor: A promising therapeutic strategy for blood malignancies. Oncology Reports, 40(1):3–15, 2018.

[23] Spencer W Hinds, Shimon A Reisner, Antonio F Amico, and Richard S Meltzer. Diagnosis of pericardial abnormalities by 2d-echo: a pathology-echocardiography correlation in 85 patients. American heart journal, 123(1):143–150, 1992.

[24] Nobuya Koike, Seung-Hee Yoo, Hung-Chung Huang, Vivek Kumar, Choogon Lee, Tae-Kyung Kim, and Joseph S Takahashi. Transcriptional architecture and chromatin landscape of the core circadian clock in mammals. Science, 338(6105):349–354, 2012.

[25] Elzbieta Kowalska, Juergen A Ripperger, Dominik C Hoegger, Pascal Bruegger, Thorsten Buch, Thomas Birchler, Anke Mueller, Urs Albrecht, Claudio Contaldo, and Steven A Brown. Nono couples the circadian clock to the cell cycle. Proceedings of the National Academy of Sciences, 110(5):1592–1599, 2013.

[26] Inna Kuperstein, E Bonnet, Hien-Anh Nguyen, David Cohen, Eric Viara, Luca Grieco, S Fourquet, Laurence Calzone, Christophe Russo, Maria Kondratova, et al. Atlas of cancer signalling network: a systems biology resource for integrative analysis of cancer data with google maps. Oncogenesis, 4(7):e160–e160, 2015.

[27] Bryan Landgraf, FE Cohen, KA Smith, R Gadski, and TL Ciardelli. Structural significance of the c-terminal amphiphilic helix of interleukin-2. Journal of Biological Chemistry, 264(2):816–822, 1989.

[28] Nicolas Le Novere, Michael Hucka, Huaiyu Mi, Stuart Moodie, Falk Schreiber, Anatoly Sorokin, Emek Demir, Katja Wegner, Mirit I Aladjem, Sarala M Wimalaratne, et al. The systems biology graphical notation. Nature biotechnology, 27(8):735–741, 2009.

[29] Takuya Matsuo, Shun Yamaguchi, Shigeru Mitsui, Aki Emi, Fukuko Shimoda, and Hitoshi Okamura. Control mechanism of the circadian clock for timing of cell division in vivo. Science, 302(5643):255–259, 2003.

[30] Huaiyu Mi, Falk Schreiber, Stuart Moodie, Tobias Czauderna, Emek Demir, Robin Haw, Augustin Luna, Nicolas Le Nov`ere, Anatoly Sorokin, and Alice Villéger. Systems biology graphical notation: activity flow language level 1 version 1.2. Journal of integrative bioinformatics, 12(2):340–381, 2015.

[31] Huaiyu Mi and Paul Thomas. Panther pathway: an ontology-based pathway database coupled with data analysis tools. In Protein Networks and Pathway Analysis, pages 123–140. Springer, 2009.

[32] Aurélien Naldi, Céline Hernandez, Nicolas Levy, Gautier Stoll, Pedro T Monteiro, Claudine Chaouiya, Tomáš Helikar, Andrei Zinovyev, Laurence Calzone, Sarah Cohen-Boulakia, et al. The colomoto interactive notebook: accessible and reproducible computational analyses for qualitative biological networks. Frontiers in physiology, 9:680, 2018.

[33] Kanae Oda, Yukiko Matsuoka, Akira Funahashi, and Hiroaki Kitano. A comprehensive pathway map of epidermal growth factor receptor signaling. Molecular systems biology, 1(1):2005–0010, 2005.

[34] Marek Ostaszewski, Stephan Gebel, Inna Kuperstein, Alexander Mazein, Andrei Zinovyev, Ugur Dogrusoz, Jan Hasenauer, Ronan MT Fleming, Nicolas Le Novere, Piotr Gawron, et al. Community-driven roadmap for integrated disease maps. Briefings in bioinformatics, 20(2):659–670, 2019.

[35] Loïc Paulevé. Pint: A static analyzer for transient dynamics of qualitative networks with ipython interface. In International Conference on Computational Methods in Systems Biology, pages 309–316. Springer, 2017.

[36] Loïc Paulevé. Reduction of qualitative models of biological networks for transient dynamics analysis. IEEE/ACM transactions on computational biology and bioinformatics, 15(4):1167–1179, 2017.

[37] Loïc Paulevé, Geoffroy Andrieux, and Heinz Koeppl. Under-approximating cut sets for reachability in large scale automata networks. In International Conference on Computer Aided Verification, pages 69–84. Springer, 2013.

[38] Loïc Paulevé, Juraj Kolčák, Thomas Chatain, and Stefan Haar. Reconciling qualitative, abstract, and scalable modeling of biological networks. Nature Communications, 11(1), 2020.

[39] Adrien Rougny, Christine Froidevaux, Laurence Calzone, and Lïc Paulevé. Qualitative dynamics semantics for sbgn process description. BMC systems biology, 10(1):1–24, 2016.

[40] Adrien Rougny, Vasundra Touré, Stuart Moodie, Irina Balaur, Tobias Czauderna, Hanna Borlinghaus, Ugur Dogrusoz, Alexander Mazein, Andreas Dräger, Michael L Blinov, et al. Systems biology graphical notation: process description language level 1 version 2.0. Journal of integrative bioinformatics, 16(2), 2019.

[41] Anton Shostak, Bianca Ruppert, Nati Ha, Philipp Bruns, Umut H Toprak, Roland Eils, Matthias Schlesner, Axel Diernfellner, and Michael Brunner. Myc/miz1-dependent gene repression inversely coordinates the circadian clock with cell cycle and proliferation. Nature communications, 7(1):1–11, 2016.

[42] Anatoly Sorokin, Nicolas Le Nov`ere, Augustin Luna, Tobias Czauderna, Emek Demir, Robin Haw, Huaiyu Mi, Stuart Moodie, Falk Schreiber, and Alice Villéger. Systems biology graphical notation: entity relationship language level 1 version 2. Journal of integrative bioinformatics, 12(2):281–339, 2015.

[43] Gabriele Sulli, Amy Rommel, Xiaojie Wang, Matthew J Kolar, Francesca Puca, Alan Saghatelian, Maksim V Plikus, Inder M Verma, and Satchidananda Panda. Pharmacological activation of reverbs is lethal in cancer and oncogene-induced senescence. Nature, 553(7688):351–355, 2018.

[44] Vasundra Touré, Andreas Dräger, Augustin Luna, Ugur Dogrusoz, and Adrien Rougny. The systems biology graphical notation: Current status and applications in systems medicine. 2020.

[45] Keziban Ünsal-Kaçmaz, Thomas E Mullen, William K Kaufmann, and Aziz Sancar. Coupling of human circadian and cell cycles by the timeless protein. Molecular and cellular biology, 25(8):3109–3116, 2005.

[46] Yongjun Wang, Douglas Kojetin, and Thomas P Burris. Anti-proliferative actions of a synthetic reverb*α*/*β* agonist in breast cancer cells. Biochemical pharmacology, 96(4):315–322, 2015.

[47] Xuan Zhao, Tsuyoshi Hirota, Xuemei Han, Han Cho, Ling-Wa Chong, Katja Lamia, Sihao Liu, Annette R Atkins, Ester Banayo, Christopher Liddle, et al. Circadian amplitude regulation via fbxw7-targeted reverb*α* degradation. Cell, 165(7):1644–1657, 2016.

[48] Xianlin Zou, Dae Wook Kim, Tetsuya Gotoh, Jingjing Liu, Jae Kyoung Kim, and Carla V Finkielstein. A systems biology approach identifies hidden regulatory connections between the circadian and cell-cycle checkpoints. Frontiers in Physiology, 11, 2020.

