## Supplementary Information for "A Detailed Map of Coupled Circadian Clock and Cell Cycle with Qualitative Dynamics Validation"

November 6, 2020

### 1 Modification of the maps for the merging

#### 1.1 Modifications to the cell cycle map

- We added the transcriptional effects of E2F1 (under different forms) on the expression of the cyclin A2, cyclin B1 and cyclin E1 genes that were shown to be necessary for the progression of the mammalian cell cycle (ref). These effects were added under the form of stimulations of processes that produce the proteins from a source rather than on the expression of the genes, because the map did not contain the genes, but did contain the proteins. We considered the following forms for E2F1: in a complex with DP1* only, and in a complex with pRB* phosphorylated one, two or three times.

- We removed the translocation of DP1* from the cytosol to the nucleus and added the translocation its translocation from the nucleus to the cytosol. We also added PP2A as a catalyzer of the dephosphorylation of DP1* in the nucleus.

The following changes were made in the naming in order to ensure that nodes shared with the CLOCK map have a unique identifier, in preparation of the merging:

- We renamed the nucleus compartment to “nucleoplasm” and the cytosol compartment to “cytoplasm”

- We renamed entity pools by replacing the following strings: “p16INK4a*” by ”p16INK4a”, ”p53*” by ”p53”, ”p21Cip*” by ”p21Cip”, ”cyclin B1*” by ”CCNB1”, ”CDC2” by ”CDK1”, ”CDK1/CCNB1” by ”CCNB1/CDK1”.

- We renamed the following state variable: ”Ser 1981” to ”Ser1981”.

- We removed all complexes having both p53 and MDM2 as subunits and all related processes, because they were better described in the CLOCK map.

- We extended the sites of the p53 protein with two undefined and unset state variables because they were present on the p53 protein of the CLOCK map.
- We replaced unphosphorylated WEE1 (at an undefined site) by the more precisely described one of the CLOCK map (having two unphosphorylated sites at Ser53 and Ser472). Analogously we removed WEE1 phosphorylated at an undefined site and added two new forms of WEE1 described in the CLOCK map: phosphorylated at Ser53 and phosphorylated at Ser472. We also added processes producing each form, and catalyzed by phosphorylated CHEK1 for the form phosphorylated at Ser472, and catalyzed by a CDK1/CCNB1 complex for the form phosphorylated at Ser53. Finally, we added the catalysis of the phosphorylation of CDK1 at Tyr15 inside the CDK1/CCNB1 complex by WEE1 phosphorylated at Ser472.

##### 1.2 Modification of the circadian clock map

- CellDesigner does not allow building complex logical functions as inputs of modulations. Hence we replaced the modulations targeting the expression of the p21cip gene by a necessary stimulation having as input a complex logical function programmatically. The necessary stimulation reads as follows: the expression of p21cip is only possible when the p21cip gene is present and either protein p53 phosphorylated at Ser15 and Ser20 is present or protein RORG is present and protein NR1D1 phosphorylated one time is absent.
- Likewise, we replaced the modulations targeting the expression of the Per2 gene by a necessary stimulation that reads as follows: the expression of Per2 is only possible when the Per2 gene is present and either the complex including GC and acetylated GCR is present, or protein HSF1 is present, or the complex including the CK2$\alpha$ protein, phosphorylated and acetylated BMAL1, and protein CLOCK is present and the big complex including protein CSNK1$\epsilon$, protein CRY1, protein CLOCK, protein PER1, phosphorylated and acetylated BMAL1 and protein CRY2 is absent.

#### 2 Alernatives for the choice of the precursor state

The computation of all possible initial states as described in section 2.3.2 resulted in 24 equivalent states in regards of the constraints we considered. Each of the 24 states included 104 entities, 100 of which were common to all states. Hence we investigated the nature of the entities that did not belong to all states, and how they made up the 24 states. As a result, we found that the 24 sets resulted from a combination of the following four alternatives:

- cytoplasmic p53 or nucleoplasmic p53

- cytoplasmic E2F4 or nucleoplasmic E2F4
- cytoplasmic DP2 or nucleoplasmic DP2
- NAD+, NAM, or NMN

The first three alternatives can be explained by the fact that in the merged map, p53, E2F4, and DP2 have two possible localizations (cytoplasm and nucleoplasm), and that it contains cycling translocation processes from one localization to the other for each of these entities. As for the fourth alternative, it can be explained by the presence in the merged map of a cycle producing and consuming the three entities NAD+, NAM and NMN.
